# Learning to Encode Cellular Responses to Systematic Perturbations with Deep Generative Models

**DOI:** 10.1101/2020.01.14.906768

**Authors:** Yifan Xue, Michael Q. Ding, Xinghua Lu

## Abstract

Components of cellular signaling systems are organized as hierarchical networks, and perturbing different components of the system often leads to transcriptomic profiles that exhibit compositional statistical patterns. Mining such patterns to investigate how cellular signals are encoded is an important problem in systems biology. Here, we investigated the capability of deep generative models (DGMs) for modeling signaling systems and learning representations for transcriptomic profiles derived from cells under diverse perturbations. Specifically, we show that the variational autoencoder and the supervised vector-quantized variational autoencoder can accurately regenerate gene expression data. Both models can learn representations that reveal the relationships between different classes of perturbagens and enable mappings between drugs and their target genes. In summary, DGMs can adequately depict how cellular signals are encoded. The resulting representations have broad applications in systems biology, such as studying the mechanism-of-action of drugs.

## Introduction

A cellular signaling system is a signal processing machine that detects changes in the internal or external environment, encodes these changes as cellular signals, and eventually transmits these signals to effectors, which adjusts cellular responses accordingly. Cellular responses to perturbations often involve changes in transcriptomic programs (Azeloglu and Iyengar, 2015; Radhakrishnan et al., 2010; Weng et al., 1999). The investigation of cellular signaling systems remains an important task in the field of systems biology. A common approach is to systematically perturb a cellular system with genetic or pharmacological perturbagens and monitor transcriptomic changes in order to reverse engineer the system and gain insights into how cellular signals are encoded and transmitted. This approach has been employed in many large-scale systems biology studies, e.g., the yeast deletion library (Giaever and Nislow, 2014), the Connectivity Map project (Lamb, 2007; Lamb et al., 2006), and its more recent iteration: the Library of Integrated Network-based Cellular Signatures (LINCS) (Keenan et al., 2018; Subramanian et al., 2017). The LINCS project is arguably the most comprehensive systematic perturbation dataset currently available, in which multiple cell lines were treated with over tens of thousands perturbagens (e.g., small molecules or single gene knockdowns), followed by monitoring gene expression profiles using a new technology known as the L1000 assay that utilizes ~1,000 (978) landmark genes to infer the entire transcriptome (Subramanian et al., 2017).

While there are numerous studies using LINCS data to investigate the mechanism-of-action (MOA) of drugs and to promote clinical translation of MOA information (Iwata et al., 2017; Lamb et al., 2006; Pabon et al., 2018; Siavelis et al., 2015; Subramanian et al., 2017; Wang et al., 2016), few studies aim to use the data to learn to represent the cellular signaling system as an information encoder and examine how different perturbagens affect the system. It can be imagined that perturbing different signaling components at different levels of a signaling cascade would lead to compositional statistical structure in the data. The responses to perturbations of an upstream signaling molecule will likely subsume the responses to perturbations of its downstream molecules. Capturing such a compositional statistical structure requires models that are capable of representing the hierarchical relationships among signaling components. In this study, we developed deep learning models, specifically deep generative models (DGMs) to understand how perturbagens affect the cellular signal encoding process and lead to changes in the gene expression profile.

DGMs are a family of deep learning models which employ a set of hierarchically organized latent variables to learn the joint distribution of a set of observed variables. After training, DGMs are capable of generating simulated data that preserve the same compositional statistical structure as the training data. The hierarchical organization of latent variables is particularly suitable for representing cellular signaling cascades and detecting compositional statistical patterns derived from perturbing different components of cellular systems. The capability to “generate” samples similar to the training data is of particular interest. If a model can accurately regenerate transcriptomic data produced under different perturbations, the model should have learned a representation of the cellular signaling system that enables it to encode responses to perturbations. Such representations would shed light on the MOAs through which perturbagens impact different cellular processes.

In this study, we implemented two DGMs, the variational autoencoder (VAE) (Figure 1A) (Hinton and Salakhutdinov, 2006; Kingma and Welling, 2014; Rezende et al., 2014) and a novel model designed by us, the supervised vector-quantized variational autoencoder (S-VQ-VAE) (Figure 1B). These models were deployed to learn latent representations of the signaling state of cells studied in the LINCS project. We show that the VAEs can reconstruct the distribution of the input data accurately and generate new data that are indistinguishable from real observed data using a relatively small number of latent variables that capture major signatures. We designed S-VQ-VAE, extended from the vector-quantized VAE (VQ-VAE) (Van Den Oord and Vinyals, 2017), for learning a unique representation for each class of perturbagens. We demonstrate that by adding a supervised learning component to VQ-VAE, we are able to summarize the common features of a family of drugs into a single embedding vector and use these vectors to reveal correlations between different families of drugs. To our knowledge, this is the first study that systematically investigates the power of DGMs for learning statistical structures of gene expression profiles, and our findings support the use of deep generative models as a powerful tool in modeling cell signaling systems.

**Figure 1.**
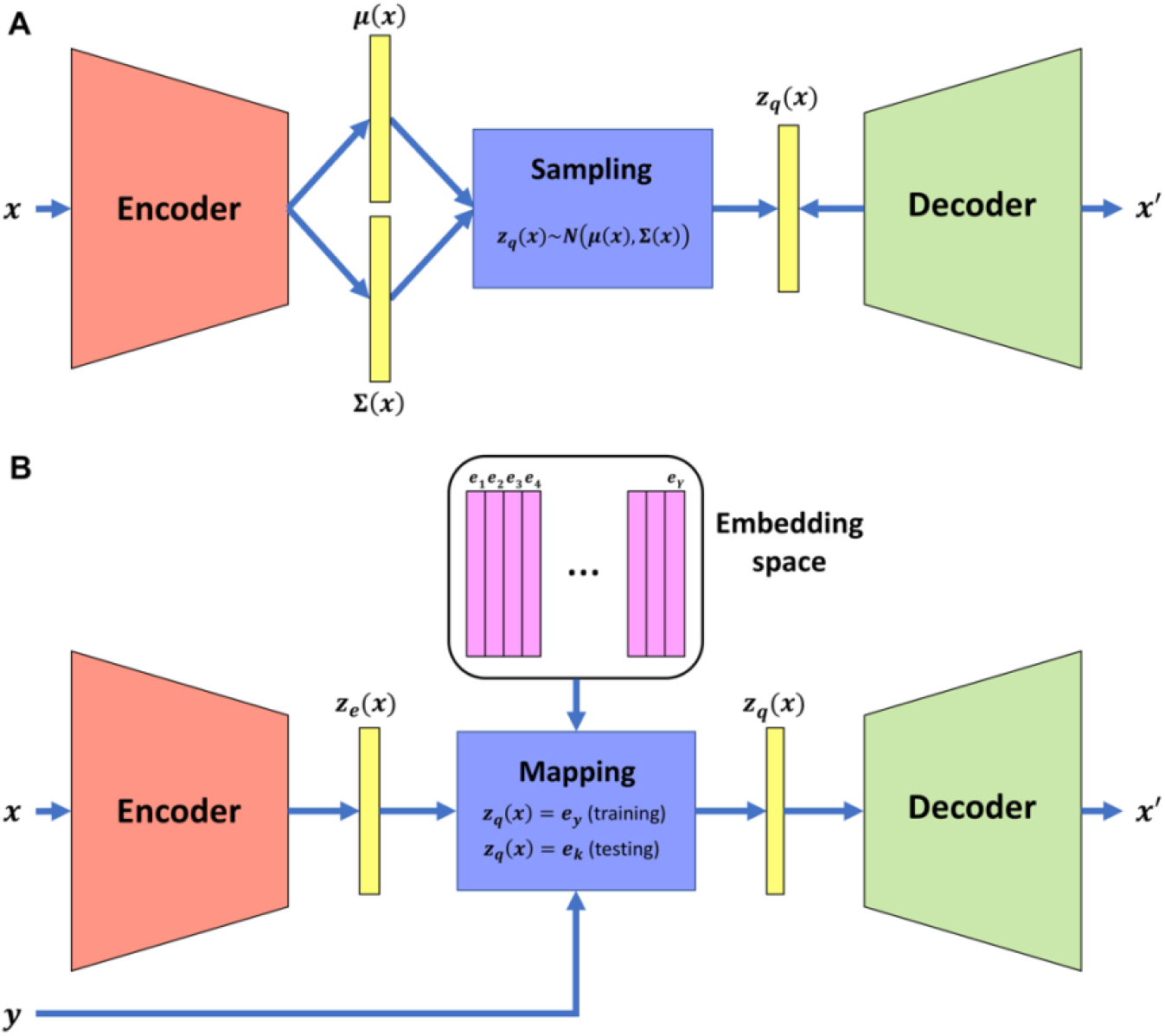
The VAE model and S-VQ-VAE model. (A) The architecture of VAE. The encoder and decoder are two sub-neural networks. An input case is transformed into a mean vector ***μ*(*x*)** and a covariance vector ***∑*(*x*)** by the encoder, from which the encoding vector ***z_q_*(*x*)** is sampled and fed to the decoder to reconstruct the input case. The distribution of the encoding vector is trained to follow a prior standard normal distribution. (B) The architecture of S-VA-VAE. S-VQ-VAE is an extension of VQ-VAE where the training of the embedding space is guided by the label of the input data. Similar to VAE, an input case is first transformed into an encoding vector ***z_e_*(*x*)** by the encoder. During training, the encoding vector is replaced by the embedding vector ***e_y_*** designated to represent the label *y* of data to reconstruct the input case. The embedding vector is updated according to the reconstruction error. During testing, the encoding vector is replaced by the nearest neighbor embedding vector ***e_k_***.

## Results

### Modeling Cellular Transcriptomic Processes with VAE

To determine the optimal VAE architecture for learning expression profiles of LINCS data, we carried out a series of model comparison experiments and selected the architecture based on model complexity, reconstruction error and other aspects of performance (see Methods). The input and output layers contained 978 nodes, each corresponding to one of the 978 landmark genes in an L1000 expression profile. The internal architecture was composed of three hidden layers in its encoder, with 1000, 1000, and 100 hidden nodes, respectively (Figure S1, each hidden node corresponds to a latent variable); the decoder had a reverse architecture as the encoder. We trained three VAE models independently on gene expression datasets consisting of different combinations of samples treated with two types of perturbagens: the first model was trained on the small molecule perturbagen (SMP) dataset that contained 85,183 expression profiles from seven cell lines treated with small molecules (Table S1); the second model was trained on the genetic perturbagen (GP) dataset that contained 116,782 expression profiles from nine cell lines with a single gene knockdown (Table S1); and the third model was trained on the combined SMP and GP dataset (SMGP). We excluded 4,649 samples treated with two proteasome inhibitors, bortezomib and MG-132, from the SMGP dataset as these samples form a unique outlier distribution on the principal component analysis (PCA) plot (Figure S2, see Methods). Table S2 shows the performance of the three VAE models trained independently on the SMP, GP, and SMGP datasets.

We first examined whether the VAE models are able to capture the distribution of the input data by generating new data and comparing the distribution of the generated data with the original input data. For each of the three models, we randomly generated 10,000 samples and projected them with 10,000 randomly selected original training samples into the first two components PCA space (Figure 2). From the scatter plots in Figure 2 (A, D and G), we can see that the VAE-generated data points take up a similar space in the PCA plot as the input data for all three datasets. The consistency in the centroid, shape, and range of the density contour indicate that the VAE models are able to reconstruct the distribution of the input data (Figure 2).

**Figure 2.**
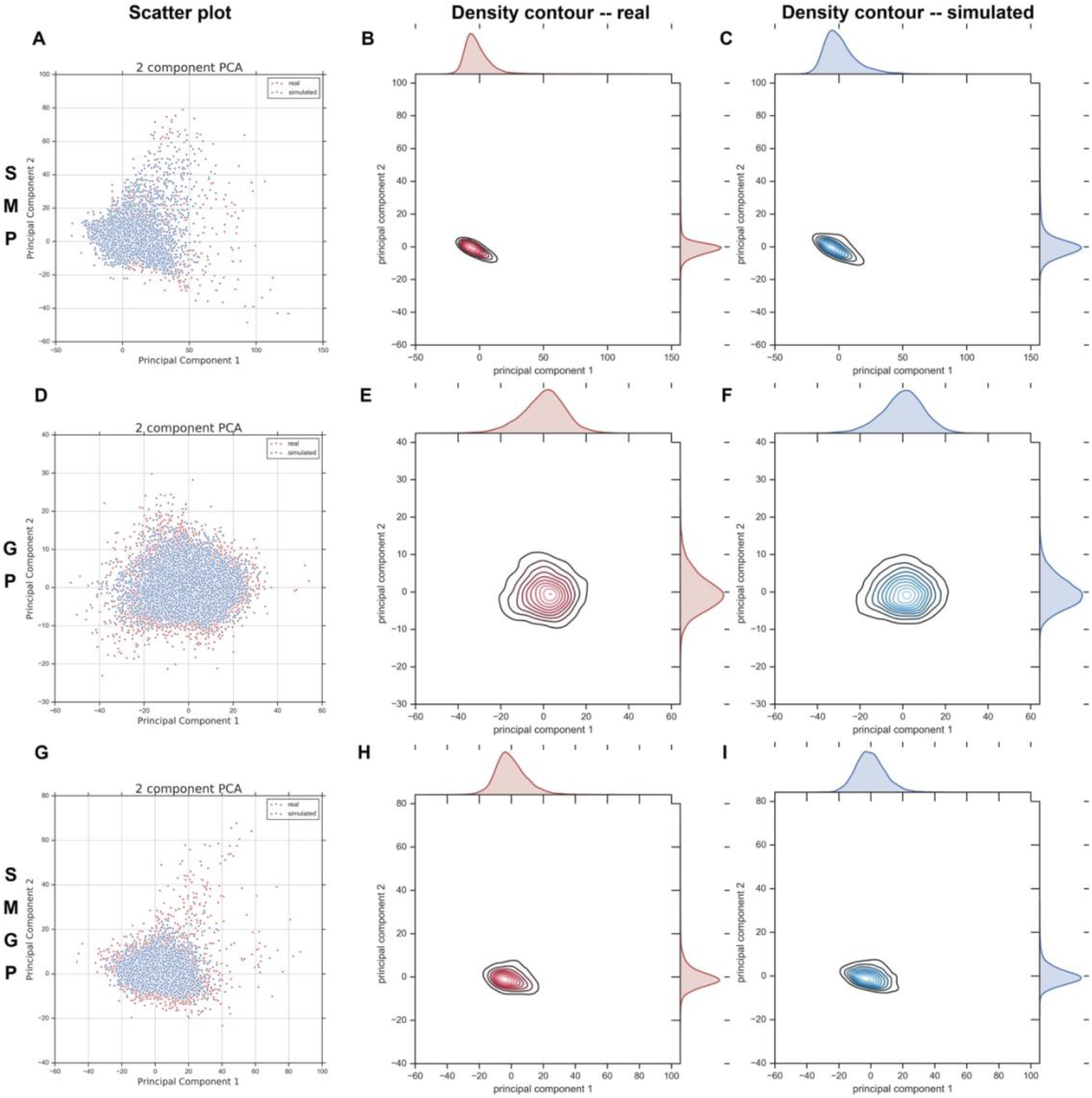
Simulated data of VAE vs. original input data. (A) Scatter plot of simulated data (blue points) generated by SMP-trained VAE and the original SMP data (red points) in the space of the first two PCA components. (B) The density contour of the real data in (A). (C) The density contour of the simulated data in (A). (D) Scatter plot of simulated data (blue points) generated by GP-trained VAE and the original GP data (red points) in the space of the first two PCA components. (E) The density contour of the real data in (D). (F) The density contour of the simulated data in (D). (G) Scatter plot of simulated data (blue points) generated by SMGP-trained VAE and the original SMGP data (red points) in the space of the first two PCA components. (H) The density contour of the real data in (G). (I) The density contour of the simulated data in (G).

Further, we performed a hierarchical clustering analysis to examine whether the newly generated data are indistinguishable from real data. Using 2,000 randomly generated samples and 2,000 randomly selected original samples, we conducted hierarchical clustering with *1-Pearson correlation* as the distance metric. We cut the dendrogram at 10 clusters and computed a mixing score (see Methods) to examine whether the generated data and original data were similarly distributed across clusters. For binary-categorical data, a mixing score is of range [0.5, 1], which gives the average proportion of data from the dominant category in each cluster. A mixing score of 0.5 indicates an even mixture of the two categories of data within all clusters, and a score of 1 indicates a clear separation between the two categories across clusters. The mixing scores obtained for 10 clusters were 0.633 for SMP-trained VAE, 0.587 for GP-trained VAE, and 0.614 for SMGP-trained VAE. Therefore, there is no strong domination of either the real data or the generated data in any cluster, and the generated data are indistinguishable from the real data according to the hierarchical clustering.

The distribution comparisons and clustering results support that as a generative model, VAE can learn the statistical structures of the LINCS expression profiles derived from diverse perturbations. This indicates that the latent variables in the learned VAE hierarchy have captured the compositional relationships of cellular responses to perturbations. Since SMGP is the largest training dataset and contains samples from both the SMP and GP datasets, all following analyses were conducted using the SMGP-trained VAE unless otherwise specified.

### A Few Signature Nodes Encode the Primary Characteristics of an Expression Profile

To gain a better understanding of how VAEs encode the distribution of diverse input data, we next examined the activation patterns of hidden nodes on different layers of the SMGP-trained VAE model. We paid particular attention to the top hidden layer of 100 nodes that serves as an “information bottleneck” for compressing the original data; this layer also serves as the starting layer for the generation of new samples. After training, each training case was associated with an instantiated encoding vector by feeding the expression profile through the encoder to the top hidden layer (*z_q_*(*x*) in Figure 1A). We noted that 12 out of 100 nodes in the encoding vector had a high variance in values across training samples (absolute average node value > 0.5, Figure 3A). For an ordinary VAE model, the prior distribution of the encoding vector is a standard normal distribution with a mean vector *μ*(*x*) = 0 and a diagonal covariance matrix *∑*(*x*) = *diag*(1) (Figure 1A), and an element of the vector shrinks towards 0 unless it is driven by data to deviate from 0. Therefore, the significantly high absolute values taken by these 12 hidden nodes suggest that they encode major signals of input data. From now we denote these 12 hidden nodes as the signature nodes.

**Figure 3.**
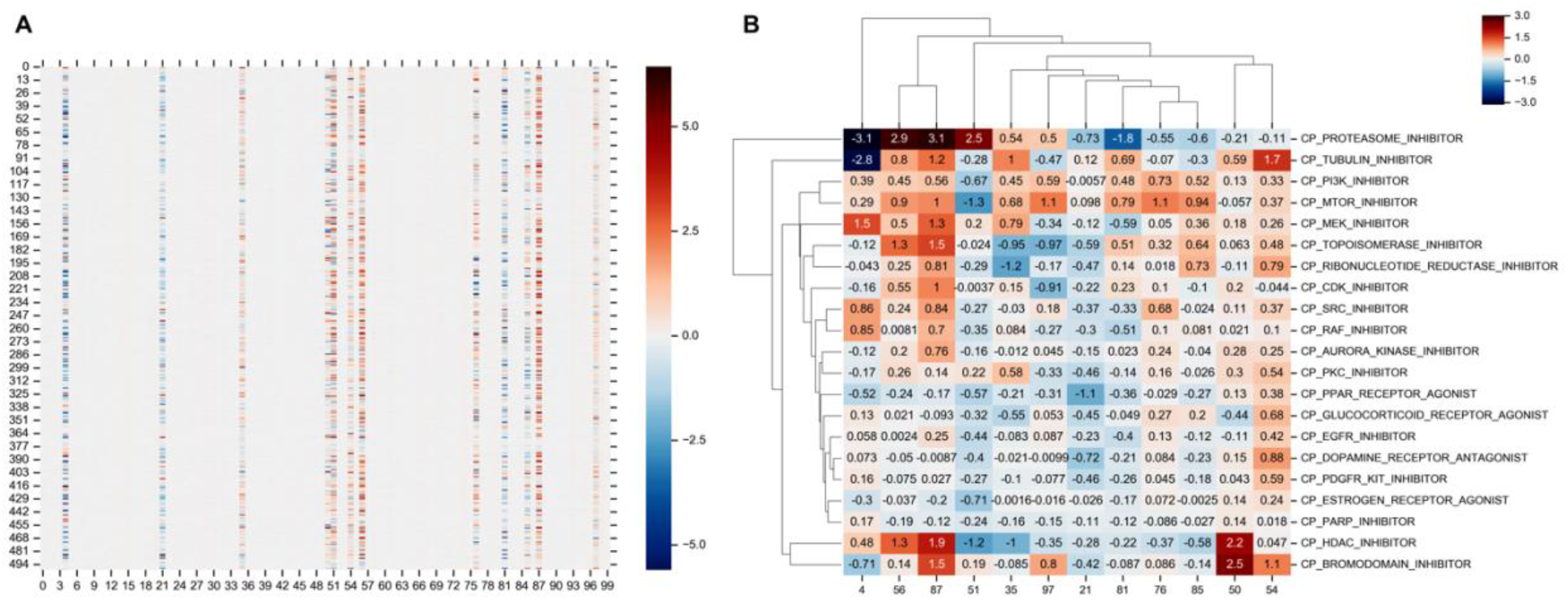
Signature nodes on the top hidden layer of SMGP-trained VAE. (A) The heatmap of the 100 hidden nodes of the top hidden layer for 500 random selected SMGP samples. The pseudo-colors represent the values of elements in the encoding vectors. (B) The average of signature nodes for samples treated with major PCLs.

We investigated whether the patterns of these 12 signature nodes reflect the MOA of drugs by examining their association with perturbagen classes (PCLs) defined by the LINCS project (Subramanian et al., 2017). We extracted a subset of the SMP dataset where the samples were treated with small molecules that had been classified into one of the PCLs based on MOA, gene targets or family, and pathway annotations (Subramanian et al., 2017). We call this subset the SMP dataset with Class information (SMC) dataset. The SMC dataset consisted of 12,079 samples treated with small molecules from 75 PCLs, and a PCL was considered as a major PCL if it had over 150 samples. We fed the samples of a major PCL through the trained VAE encoder and took the average signature node values as a representation of the PCL. As shown in Figure 3B, different PCLs presented different patterns in the signature nodes.

We further examined whether the representations of each PCL revealed correlations between PCLs via hierarchical clustering analysis (Figure 3B). PCLs that showed similarities in the signature node pattern and were closely clustered tend to share similar MOAs (Figure 3B). For example, the mTOR inhibitor and PI3K inhibitor were grouped together according to their consistent activation directions (positive vs. negative) for most signature nodes, and they are both known to impact the PI3K/AKT signaling pathway (O’Reilly et al., 2006), where the mTOR, designating the mammalian target of rapamycin (a serine/threonine kinase), is a downstream effector of PI3K. Other examples include the grouping of Src inhibitor and Raf inhibitor, where Src is known to activate Ras-c, which in turn activates Raf in the Raf-MEK-ERK pathway (Moon et al., 2002); the grouping of topoisomerase inhibitor and ribonucleotide reductase inhibitor, where both impact the DNA replication process; and the grouping of Aurora kinase inhibitor and PKC inhibitor, where Aurora kinases are essential to mediate PKC-MAPK signal to NF-κB/AP-1 pathway (Noh et al., 2015). These observations support that the 12 signature nodes preserve the crucial information of an expression profile resulting from a cellular system perturbed by small molecules.

To further demonstrate that the primary characteristics of an expression profile are encoded in the 12 signature nodes, we generated new expression profiles simulating those from cells treated with a target PCL by manipulating values of the signature nodes to mimic the pattern of the target PCL. Concretely, we preset the signature nodes to values similar to the average values of training samples treated with the target PCL as shown in Figure 3B and randomly initialized the other hidden nodes from a standard normal distribution. In this way, we randomly generated 500 new samples and compared them to real samples to see whether their nearest neighbors were from the target PCL (Figure 4A). The signature node patterns used to generate samples and the similarity of these samples to real samples and associated PCLs are shown in Figure 4B-I. In most cases, more than half of the generated data were mapped to the correct target PCL. For proteasome inhibitor and tubulin inhibitor specifically (Figure 4G and I), 100% of the generated data were nearest neighbors of real samples from the same PCL, which was repeatedly observed across independent runs. This agrees with the PCL clustering outcome in Figure 3B, where proteasome inhibitor and tubulin inhibitor were found as outliers from the other PCLs with their distinct signature node patterns. We also noted that the specific value of each signature node did not matter as long as the value correctly reflects the direction, i.e., positive or negative, of the node for a given PCL. This suggests that the major characteristics of a PCL can potentially be encoded into only 12 bits of information. The only pattern that did not have over half of the generated samples mapped to the target PCL was the mTOR inhibitor (Figure 4E). Most of the samples generated using mTOR signature nodes were closest neighbors with a PI3K inhibitor treated sample. This is reasonable, as mTOR inhibitors act downstream on the same pathway of PI3K inhibitors, thus the former’s effects can be in many cases replicated by the latter. This observation also supports that each signature node pattern reflects a specific cellular signaling process, which, after decoding, generates an expression profile that reflects how the signaling is perturbed.

**Figure 4.**
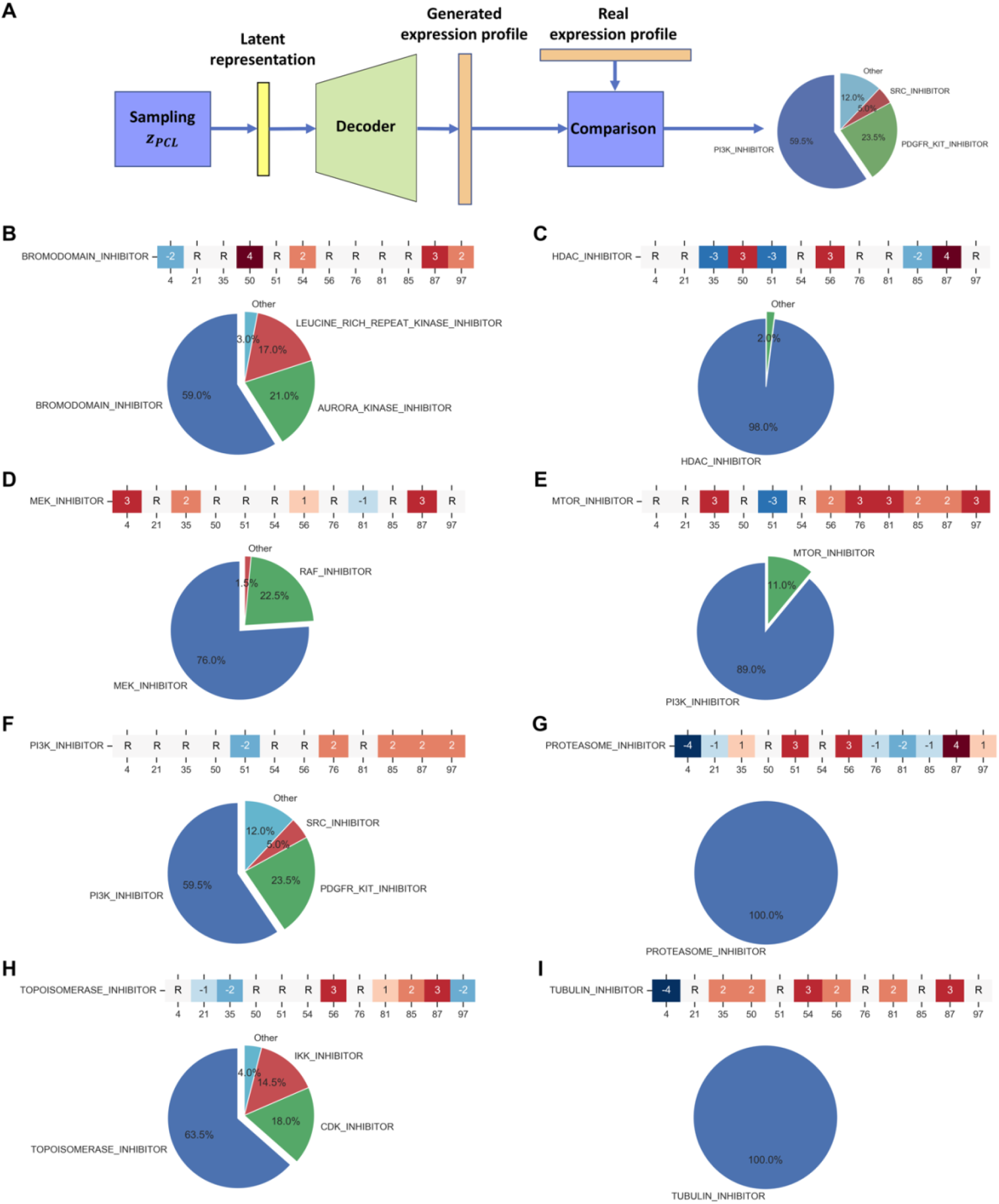
Comparison of data generated based on the signature pattern of PCLs with real data. (A) Diagram illustrating the procedure for generating new data from the signature pattern of a PCL. First, an encoding vector is initialized where the signature nodes are set according to the signature pattern of real samples from the given PCL; the non-signature nodes are randomly initialized by sampling from a standard normal distribution. The vector is then fed through the decoder of the SMGP-trained VAE, and a new expression profile is generated. The new data are compared to real data by computing the nearest neighbor based on Euclidean distance, to see if the new data are closely related to real data of the given PCL. (B-I) The composition of real data nearest neighbors of new data generated from latent representations simulating different PCLs. “R” indicates the value of the signature node is not specified but random initialized as non-signature nodes.

### Learning Global Representations of PCLs with S-VQ-VAE

The signature node representations of PCLs discussed above were obtained by averaging over samples treated with small-molecule perturbagens belonging to a PCL. In order to learn a unique, stable global representation for each PCL, we designed another DGM, the S-VQ-VAE, which utilizes the PCL label of small molecules that treated the cells to partially supervise the training process. S-VQ-VAE was extended from VQ-VAE by utilizing the vector-quantized (VQ) technique to discretize the encoding vector space into multiple mutually exclusive subspaces represented by a limited number of embedding vectors and projecting data from each class into its pre-assigned subspace (Figure 1B, see Methods). After training, each embedding vector learns to summarize the global characteristics of a class of data. In this study, we used S-VQ-VAE to learn an embedding vector with a dimension of 1000 for representing each of the 75 PCLs in the SMC dataset (Table S2).

We utilized the embedding vectors to reveal similarity and potential functional relationships between PCLs by comparing each PCL to all the others to identify its nearest neighbor based on the Pearson correlation. The nearest neighbor relationships between PCLs are visualized as a directed graph (Figure 5), in which a directed edge indicates that the source node is the nearest partner to the target node. We also used the Louvain algorithm (Blondel et al., 2008) to detect the community (aka, clusters) among PCLs, and members of different communities are indicated as pseudo-colors (Figure 5). The modularity score of the communities is 0.875, which indicates significantly denser connections existing between members within communities compared to a randomly assigned network of the same set of PCLs. Some strong relationships, like bi-directional connections, were observed in Figure 5, and many of such relationships correspond to well-documented shared MOAs between the drugs in the connected PCLs. These include the relationships that have also been revealed with the signature node representations above, e.g., the functional similarity between mTOR inhibitors and PI3K inhibitors (O’Reilly et al., 2006), and the relationship between MEK inhibitors and Src inhibitors, and Raf inhibitors (Moon et al., 2002). Other strong connections were observed between CDK inhibitors and topoisomerase inhibitors, which may reflect coordinated response to mitosis inhibition and DNA damage induction (Peyressatre et al., 2015; Weinberg, 2013), between Aurora kinase inhibitors and HDAC inhibitors that impact the histone deacetylase pathway (Li et al., 2006), and between gamma-secretase inhibitors, serotonin receptor antagonists and bile acid that affect amyloid precursor protein processing and lipid metabolism (Pimenova et al., 2014; Watanabe et al., 2010).

**Figure 5.**
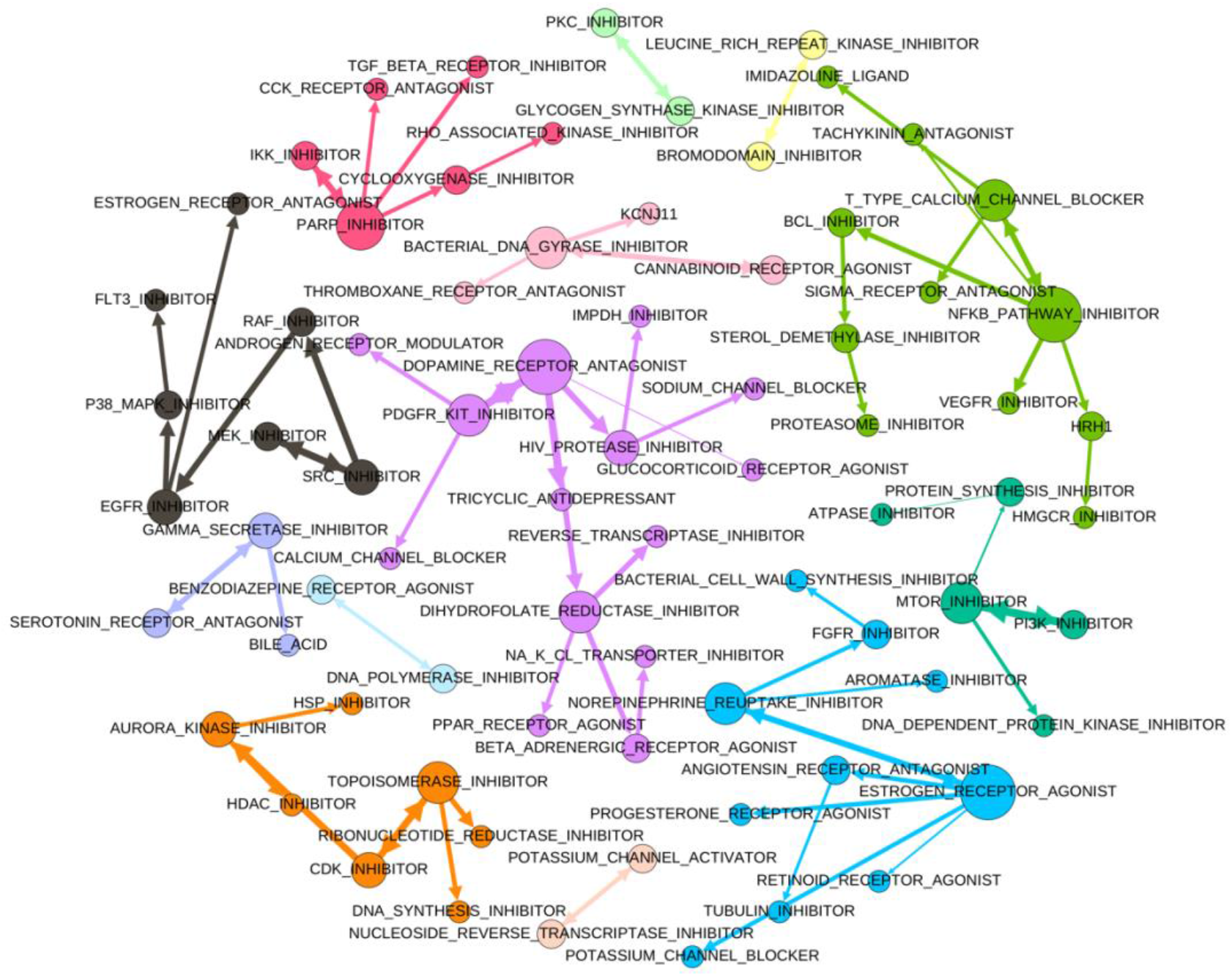
Similarity between PCLs revealed with global PCL representations learned by S-VQ-VAE. A directed edge in the graph indicates that the source node is the nearest node to the target code based on the Pearson correlation between the corresponding representations. The node size is proportional to the out-degree. The edge width is proportional to the correlation. The color of a node indicates the community the node belongs to.

The members of a PCL community also shed light on the high-level functional theme of the community. For example, the black community on the left of Figure 5 with Raf, Src, MEK, and EGFR related PCLs may represent the drug effects transmitted through the EGFR-RAS-RAF-MEK signaling cascade. The orange community (bottom left of Figure 5), consisting of inhibitors of Aurora kinase, HDAC, CDK, topoisomerase, ribonucleotide reductase, and DNA synthesis, may represent the signaling transduction for regulating DNA duplication and mitosis. The blue community (bottom right of Figure 5), with estrogen, progesterone, norepinephrine, and angiotensin may represent the comprehensive effects of perturbing hormones. These findings indicate that the global representations learned with S-VQ-VAE preserve the crucial information to reveal the functional impact of different PCLs.

### The VAE Latent Representations Preserve PCL-related Information

The latent variables at the different levels of the hierarchy of a DGM may encode cellular signals with different degrees of complexity and abstraction (Chen et al., 2016). Therefore, we next investigated the information preserved in the latent variables of different hidden layers of the SMGP-trained VAE. To do this, we first represented the SMC samples with seven types of representations, including the raw expression profiles, the latent representations obtained from the five hidden layers of the VAE (across the encoder and decoder), and the 12 signature node values (see Methods). We then used these representations to predict the PCL label of the small molecule used to treat each sample by training two multi-classification models, the logistic regression (LR) and the support vector machine (SVM). As shown in Table S3, the highest test prediction accuracy was achieved by using the raw expression profiles as input data for both LR and SVM (0.5922 and 0.5273 respectively). This was followed by the latent representations of samples extracted from the first hidden layer of the VAE encoder (0.5096 for LR and 0.4528 for SVM). The lowest accuracy was obtained using the 12 signature node values as input data (0.3814 for LR and 0.3615 for SVM). Nonetheless, the highest test accuracy achieved with latent representation, 0.5096, was nearly 10 times higher than guessing at random from the 75 unevenly distributed PCLs, 0.0543. These results indicate that although there was information loss with respect to the classification task as the representations become more abstract with deeper hidden layers, the latent representations preserved significant information from the original input data.

### The VAE Latent Representations Enhance Drug-Target Identification

Combining SMP and GP data can help establish connections between the MOAs of small molecules and genetic perturbations, which further help reveal the targets of small molecules (Lamb, 2007; Pabon et al., 2018). A simple approach is to examine whether a pair of perturbagens (a small molecule and a genetic perturbation) leads to similar transcriptomic profiles, or more intriguingly, similar latent representations that reflect the state of the cellular system (Figure 6A). Given a known pair of a drug and its target protein (Pabon et al., 2018) (Table S4), we assumed that treatment with the drug and knockdown of the gene of the protein would result in a similar transcriptomic response reflected as raw data or VAE-derived latent representations. Based on this assumption, we computed the Pearson correlations between the representation of an SMP sample (either in the form of the raw expression profile or a latent representation) and the corresponding representations of all GP samples. From these correlations, we created a ranked list of target genes knocked down in the GP samples, in a manner similar to an information retrieval task. We then compared different types of representations to identify which were more effective (measured as top and mean ranking, see Methods) for retrieving target genes than others.

**Figure 6.**
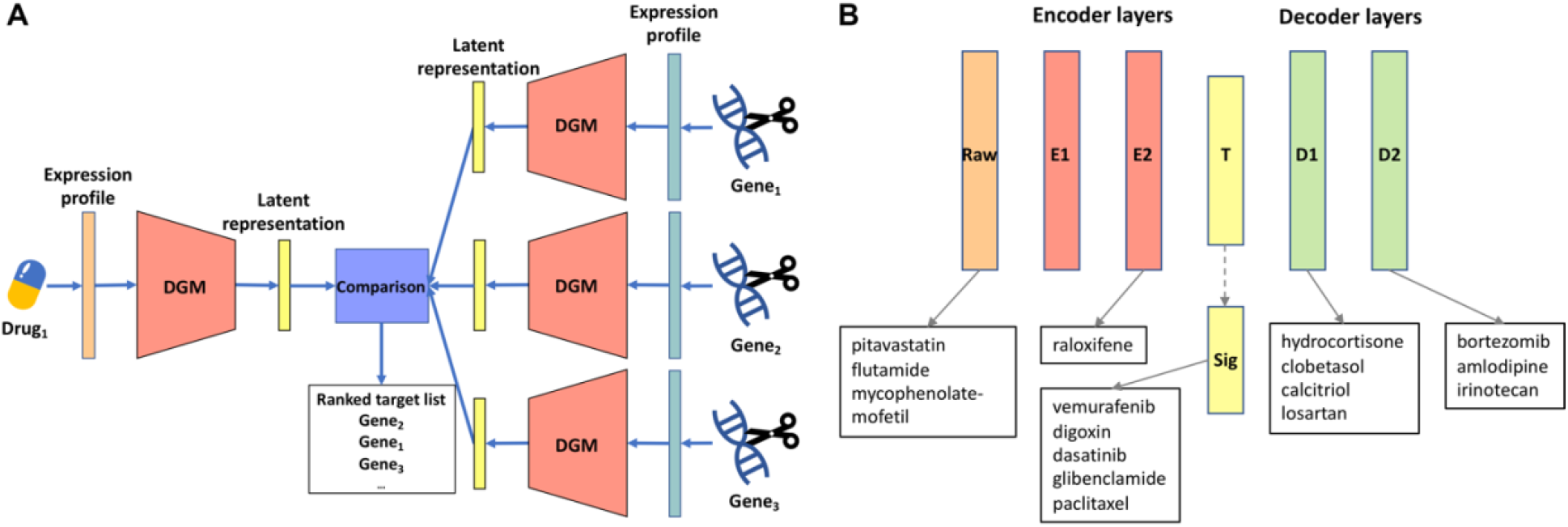
Drug-target prediction with different sample representations for 16 FDA-approved drugs. (A) Diagram illustrating the approach for drug-target prediction. For a given drug, samples treated with the drug are fed to the SMGP-trained VAE to obtain latent representations from different layers of the encoder and decoder. All GP samples are also fed to the VAE to obtain corresponding latent representations and compared with the SMP samples by computing the Pearson correlation. For a given type of representation, genes are ranked according to the correlations with respect to the representation of drug, and the ranks of the top known target of the drug are recorded. (B) The representation type that achieved the best matching (lowest mean rank) of the top known target gene for each drug.

As shown in Table S4 and Table 1, it is interesting to note that for different drugs, different representations achieved the best target-retrieval performance. This suggests that VAEs can encode the impact of different drugs with different layers in the hierarchy that potentially reflect the relative level of drug-target interactions in the cellular signaling network. Figure 6B summarizes the mean rank results where each drug is assigned to the representation layer that produced the best mean rank of its top known target(s). Most drugs have their best performance achieved with VAE-learned latent representations rather than the raw expression profiles, and for five drugs, the best performance was achieved with the 12-signature-node-representation. Table 1 gives the best rank of the top known target for each drug, which is comparable to the Table 1 in (Pabon et al., 2018). Even though our approach is essentially an unsupervised learning method based purely on expression data, 13 out of 16 drugs received an equal or better rank than that from the random forest model trained with a combination of expression and protein-protein interaction features (Pabon et al., 2018) (bolded in Table 1).

**Table 1.**
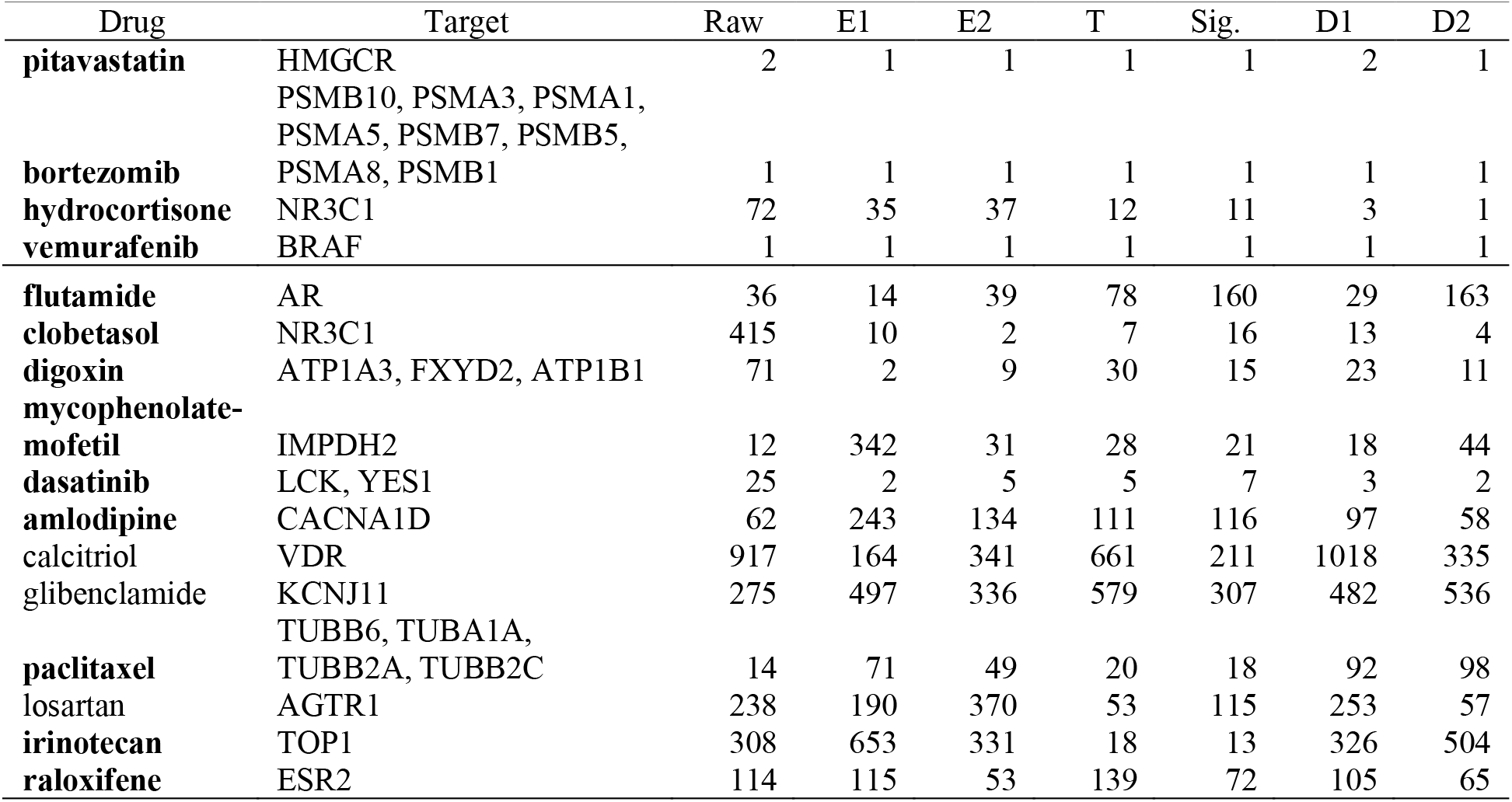
The rank of the top known target for 16 FDA-approved drugs from drug-target prediction with different types of representations. The value for a given drug and a type of representation is the top rank of the drug target gene(s) among all genes when retrieved and ranked according to the similarity between the representation of drug and representations of samples with gene knockdowns. A lower rank is better. Drugs with the lowest rank equal to or lower than the rank reported by (Pabon et al., 2018) are bolded. E: encoder layer; T: top hidden layer; Sig: signature nodes; D: decoder layer.

## Discussion

In this study, we examined the utility of DGMs, specifically VAE and S-VQ-VAE, for learning representations of the cellular states of cells treated with different perturbagens in the LINCS project. We showed that the trained VAE models are able to accurately regenerate transcriptomic profiles almost indistinguishable from the original training data. The S-VQ-VAE models are able to learn global representations from a group of transcriptomic profiles that reveal MOA associations between PCLs. These results suggest that the DGMs examined in this study are capable of learning how information is encoded in the cellular signaling system in response to diverse perturbagens. Our analyses of the DGMs also suggest that the perturbagen information can be effectively abstracted and compressed into a relatively small number of latent variables organized in a hierarchy. In particular, the encoding vectors of the VAE capture the primary characteristics of an expression profile in 12 signature nodes. Manipulating these signature nodes was sufficient to generate new data that are similar to the real data of a PCL with the same signature node pattern. Therefore, our study provides the first evidence that DGMs can be used to model the cellular signal encoding system, i.e., how cellular signals are encoded in cells. From an information theory point of view, DGMs are capable of distilling the compositional statistical structure embedded in the raw data and can use its hierarchically organized latent variables to encode the signals underlying the statistical structures in order to regenerate data.

Given the above capability of the DGMs, one interesting question is whether the latent variables in a DGM encode the signals of real biological entities, or whether they can be mapped to the signals of individual or of a set of signaling molecules. We conjecture that from an information theory point of view, this is likely the case. Instead of inventing a new coding system, the most efficient approach for a DGM to regenerate the data is to mimic how signals are encoded in real cells. It is possible to examine the cellular signals encoded by signaling molecules and compare them to the signals encoded by latent variables in a DGM, and therefore the signaling molecules may be matched to certain latent variables. This approach is supported by a previous study of the yeast transcriptomic system (Chen et al., 2016), in which a nearly one-to-one mapping can be identified between certain latent variables and yeast transcription factors. Our current results provide additional supporting evidence: latent variables in a certain layer of the DGMs were more effective in establishing connections between a given small molecule perturbagen and its known target genes than other layers, suggesting these latent variables encode the impact of the perturbagen better than others and can be potentially mapped to biological entities perturbed by the perturbagen in cells.

An explicit interpretation of the roles of latent variables in a general deep learning model is usually very hard, due to the complexity and flexibility of the model topology. Nonetheless, by adding the PCL labels as side information, we were able to obtain a clear one-to-one mapping between S-VQ-VAE embedding vectors and PCLs. This exploration suggests that if additional auxiliary information is utilized for training DGMs, it is possible to derive interpretable representations of cellular systems. Many biological sources of big data contain diverse information that can be used in such a fashion. For example, The Cancer Genome Atlas (TCGA) dataset provides genomic, epigenomic, and transcriptomic data of cancer cells resulting from genomic perturbation by nature, which can be used to model how genomic perturbations lead to transcriptomic changes in the onset of disease (Cai et al., 2018; Tao et al., 2019). We anticipate that by incorporating multiple types of data into DGMs, it will be possible to model the cellular signaling systems with enhanced accuracy and learn interpretable cellular state representations. Such accurate and interpretable representations of cellular systems would have a significant impact across biomedical research fields like drug development and precision medicine.

## Acknowledgements

This work was partially supported by the National Library of Medicine, Grant No. R01LM012011, by the National Human Genome Research Institute, Grant No. U54HG008540 via the trans-NIH Big Data to Knowledge (BD2K) Initiative (http://www.bd2k.nih.gov), and by the Pennsylvania Department of Health, Grant No. 4100070287. The content is solely the responsibility of the authors and does not necessarily represent the official views of the above funding agents.

## Author Contributions

Conceptualization, Y.X. and X.L.; Methodology, Y.X., M.D., and X.L.; Software, Y.X.; Formal Analysis, Y.X.; Investigation, Y.X.; Resources, X.L.; Writing – Original Draft, Y.X.; Writing – Review & Editing, X.L. and M.D.; Supervision, X.L.; Funding Acquisition, X.L.

## STAR Methods

### Lead Contact and Materials Availability

Further information and requests for resources should be directed to and will be fulfilled by the Lead Contact, Dr. Xinghua Lu. This study did not generate new unique reagents.

### Method Details

#### Data

The SMP dataset was extracted from the Gene Expression Omnibus (GEO) dataset GSE70138, which contained the level 5 L1000 expression data (moderate z-scores) of the 978 landmark genes of 85,183 samples from seven major cell lines treated with small molecules (Table S1). The GP dataset was from the GEO dataset GSE106127, which contained the level 5 data of 116,782 samples from nine major cell lines with gene knockdowns (Table S1). A cell line was considered as a major cell line if the cell line had over 10,000 samples. We performed PCA on the two datasets, and the distributions of samples in the first two principal components are shown in Figure S2. By comparing the scatter plot of the SMP dataset with its density contour (Figure S2 A and B), we can see that the group of samples on the right of the scatter plot is an outlier group with high variance but low density. This group contained 4,649 samples treated with two proteasome inhibitors, bortezomib, and MG-132. Therefore, in the third dataset, the SMGP dataset that merges the SMP dataset with the GP dataset, these outlier samples were excluded. The removal of outliers resulted in comparable distributions between SMP samples and GP samples (Figure S2 E and F), which enabled the use of the SMGP dataset for training a VAE model to reveal connections between small molecules and knocked down genes.

The SMC dataset was a subset of the SMP dataset that contained 12,079 samples treated with 204 small molecules belonging to 75 PCLs. The SMC dataset was used to train LR and SVM models for predicting PCLs of samples based on cellular representations learned from VAEs.

The dataset used to learn PCL representations with S-VQ-VAE was a subset of the SMC dataset, the SMCNP dataset (Table S2), which excluded the samples treated with the proteasome inhibitor MG-132 (bortezomib was not given a PCL label, and thus had been excluded from the SMC dataset). This subset contained 9,769 samples treated with small molecules from 75 PCLs.

#### S-VQ-VAE Model

S-VQ-VAE is a new DGM designed in this study for learning a vector representation (embedding) for each PCL. The model was extended from the standard VQ-VAE (Van Den Oord and Vinyals, 2017) by adding a supervised mapping step to guide the training of the embedding space. Like VQ-VAE, a S-VQ-VAE is composed of three parts, an encoder neural network to generate the encoding vector *z_e_*(*x*) given an input vector *x*, an embedding space to look up the discrete representation *z_q_*(*x*) based on *z_e_*(*x*), and a decoder neural network to reconstruct the input data from *z_q_*(*x*) (Figure 1B). Suppose that the encoder encodes the input data to a vector of length *D*, the embedding space *E* is then defined as *E∈R_YxD_*, where *Y* is the number of different classes of the input data. In our case, this corresponds to PCLs. Each of the *Y* embedding vectors of dimension *D* is designated to learn a global representation of one of the classes. In forward computation, an input *x* is first converted to its encoding vector *z_e_*(*x*), which will be used to update the embedding space. In the training phase, *z_e_*(*x*) is replaced with *z_q_*(*x*)=*e_y_* to pass to the decoder, where *e_y_* is the embedding vectors of the class *y* of *x*. In the testing phase, *z_e_*(*x*) is replaced by its nearest code *z_q_*(*x*)=*e_k_* with

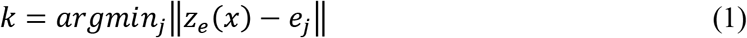

Note that we are not assuming a uniform distribution of the embedding vectors as in the ordinary VQ-VAE (Van Den Oord and Vinyals, 2017). Instead, the distribution of codes is determined by the input data with its discrete class labeling governing by a multinomial distribution.

In order to design a model that can learn individual representations through data reconstruction as well as learn a global representation for each class in a supervised manner, the objective function of S-VQ-VAE contains a reconstruction loss to optimize the encoder and decoder (first term in Equation 2), and a dictionary learning loss to update the embedding space (second term in Equation 2). The form of reconstruction loss can be selected based on the data type, and here we used the mean square error (MSE). Following the training protocol of standard VQ-VAE (Van Den Oord and Vinyals, 2017), we chose VQ as the dictionary learning algorithm, which computes the *l_2_* error between *z_e_*(*x*) and *e_y_* thus updating the embedding vector towards the encoding vector of an input case of class *y*. To control the volume of the embedding space, we also added a commitment loss between *z_e_*(*x*) and *e_y_* to force the individual encoding vector towards to the corresponding global embedding vector (third term in Equation 2).

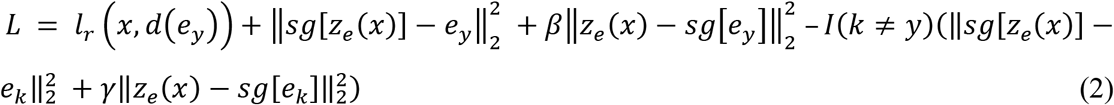

In addition to making the encoding vectors and the embedding vectors converge, we added two additional terms to force the encoding vector of an input data to deviate from the nearest embedding vector *e_k_* if *k≠y* (i.e., to minimize misclassification with the nearest neighbor). As given in Equation 2, the fourth term is another VQ objective which updates the embedding vector of the mis-class. The final term, called the divergence loss, expands the volume of the embedding space in order to allow different classes to diverge from each other. Coefficients are applied to the commitment loss (*β*) and divergence loss (*γ*) to control the strength of regularization over the embedding space volume. According to preliminary experiments using coefficients from [0, 1], the performance of the model is quite robust to these coefficients. For generating the results presented in this study, we used *β* = 0.25, and *γ* = 0.1. Note that the mapping step with either the class label or nearest neighbor has no gradient defined for it. As in training VQ-VAE, we approximate the gradient in a manner similar to the straight-through estimator (Bengio et al., 2013), by passing the gradient from the reconstruction loss from *z_q_*(*x*) directly to *z_e_*(*x*).

As a generative model, S-VQ-VAE can also be used to generate new data from the distribution of the training data. The data generation process is composed of two steps, similar to the ancestral sampling method. First, sample a target class *y* from the distribution of classes of the input data. Second, sample an encoding vector *z ~ N*(*e_y_, σ_2_*), where σ_2_ is the covariance matrix of hidden variables estimated from the training data of class *y*. A new sample of class *y* can then be generated by passing *z* to the decoder of S-VQ-VAE. The generation process reflects another advantage of S-VQ-VAE compared to unsupervised GMs: we can determine what content the new data should present rather than interpret it afterward.

In this study, we only utilized the global representation learning function of S-VQ-VAE. The test phase and the new data generation function of S-VQ-VAE were not examined here. To see how S-VQ-VAE can be used as a general generative model, please refer to our tutorial of S-VQ-VAE at https://github.com/evasnow1992/S-VQ-VAE, where we provide an example applying S-VQ-VAE on a benchmark machine learning dataset, the MNIST handwritten digits data (LeCun et al., 1998).

#### Model Architecture and Training Setting

The VAE model we implemented had three hidden layers in its encoder and three hidden layers in its decoder; the third hidden layer of the encoder was shared by both the encoder and decoder parts via a sampling step (Figure S1) and is also called the top hidden layer. The structure of the encoder was 978-1000-1000-100, where we had 978 nodes in the input layer, each corresponding to a landmark gene in the LINCS data, 1000 nodes in the first and second hidden layers, and 100 nodes in the third hidden layer (Figure S1). The structure of the decoder was just the reverse of the encoder. We only included the 978 landmark genes as input data to avoid redundant information from the inferred expression levels of other genes. The number of hidden layers and the number of nodes on each layer were determined based on preliminary experiments with a wide range of model architectures. Specifically, we tried architectures from 978-500-15 to 978-2000-1000-200 to select a model with as a simple structure as possible and with a low training error. Based on our previous experience, a three hidden layer model with 1000-1500 nodes on the first hidden layer, ~1000 nodes on the second hidden layer and small bottleneck on the third hidden layer usually performs the best (Chen et al., 2016; Ding et al., 2018). The best model we achieved in this study had a structure of 978-1000-1000-100.

We used a standard normal distribution, *N*(0, 1), as the prior distribution of the top hidden layer variables *p*(*z*) of VAE. The input data of our models were the L1000 level 5 gene expression data of range [-10, +10]. In order to preserve the sign information of the input data, where a positive value indicates high-expression of a gene and a negative value indicates low-expression of a gene, we chose the tangent function as the activation function for all hidden layers. Note that the tangent function will map a real number to [-1, +1], while our input data are of range [-10, +10]. In order to reconstruct the input data, the outputs of the last layer of the decoder were rescaled to [-10, +10] before computing the reconstruction loss (Figure S1).

The loss/target function for training a general VAE is

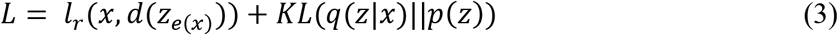

where the first term is the reconstruction loss and the second term is the KL-distance between the posterior distribution of the top hidden variables *q*(*z*|*x*) given the input data and the prior variational distribution *p*(*z*). In our implementation, we computed the MSE as the reconstruction loss. We trained three VAE models using the SMP, GP, and SMGP datasets independently. Each model was trained on 9/10 (random split) of the data and validated on the other 1/10 data. All models were trained for 300 epochs, with batch size 512 and learning rate 1e-3 (Table S2). To generate a new sample, we first sampled from the multi-variate *N*(0, 1) distribution to get an encoding vector, then passed the vector through the decoder of the VAE to generate a new data.

The S-VQ-VAE model we implemented had a single hidden layer of 1,000 nodes in its encoder. The decoder was the reverse of the encoder. As in VAE, we also used the tangent activation function for S-VQ-VAE and rescaled the data from [-1, 1] to [-10, 10] before computing the reconstruction loss. The number of hidden layers and hidden nodes were selected based on preliminary experiments with architectures from one to two hidden layers and 200 to 1500 hidden nodes in each layer. The embedding space contained 75 codes, one for each PCL. The model was trained on 9/10 (random split) of the SMCNP dataset for 900 epochs, with batch size 256, and learning rate 1e-4. The model was validated on the other 1/10 data (Table S2).

#### PCL Prediction

Seven different types of sample representations were evaluated as predictors for modeling the PCL label of the small molecule that perturbed each SMC sample via LR and SVM. The representations included the raw expression profile, the latent representations from three encoder layers of the SMGP-trained VAE, the 12 signature nodes values, and the latent representations from two decoder layers of the SMGP-trained VAE (the top hidden layer of the encoder is shared with the decoder, thus there are only two independent decoder layers). The latent representation of a layer of a sample was obtained by feeding the expression profile of the sample to the pre-trained VAE and extracting the values of hidden nodes on the desired layer. The prediction accuracy reported in this study was obtained by doing a 10-fold cross-validation across SMC data. Specifically, the SMC data were randomly split into 10 subsets. In each iteration, an independent model was trained on 9 of the subsets and validated on the 10th subset. The reported accuracy is the average validation accuracies over the 10 models.

#### Drug-Target Prediction

The known drug-target relationships were extracted from the ChEMBL database (Gaulton et al., 2011) referring to the Table 1 of (Pabon et al., 2018), which included 16 drugs tested in all the seven major cell lines in the SMP dataset. Here we considered different LINCS drug IDs with the same drug name as the same perturbagen. For predicting the gene targets for each drug, we first extracted samples treated with the drug from the SMP dataset. Then for each sample, we computed the Pearson correlations between the representation of the sample and the corresponding representations of all 116,782 samples from the GP dataset. The genes knocked down in the GP samples were ranked according to the Pearson correlations, and the rank of the top known target gene was recorded. Finally, the best top rank and mean top rank across all samples treated with the same drug were computed and used to compare different types of representations. Similar to PCL classification, seven types of sample representations were compared based on the top rank and mean rank.

### Quantification and Statistical Analysis

#### Program Language, Packages, and Softwares

VAE and S-VQ-VAE models were implemented in Python using the library *PyTorch* 0.4.1 (Paszke et al., 2017). Adam optimizer was used for updating the models. PCA analysis, LR functions, and SVM functions were from the Python library *Scikit-learn* 0.21.3 (Pedregosa et al., 2011). For LR, we used random seed 0 for shuffling data and solver “lbfgs” (Limited-memory BFGS) for multi-classification. For SVM we used random seed 0 and default settings for the other hyper-parameters. Distance computation functions, including Euclidean distance and Pearson correlation, related to Figure 3B, Figure 4, Figure 5, and drug-target prediction were from the Python library *SciPy* 1.3.1 (Virtanen et al., 2019). For Figure 3B and Figure 4 we used the Euclidean distance for revealing general associations between expression profile representations and for Figure 5 and drug-target prediction we used the Pearson correlation for emphasizing more on the orientation consistency between representations. Hierarchical clustering and heatmap visualization related to Figure 3 were carried out with the Python library *Seaborn* 0.9.0 (Waskom, 2018). S-VQ-VAE PCL representation graph visualization and community detection related to Figure 5 were accomplished with software *Gephi* 0.9.2 (Bastian et al., 2009). The community detection algorithm being used was the Louvain algorithm developed by (Blondel et al., 2008) and was run with randomization (for better decomposition), using edge weights, and resolution 1 (for detecting smaller communities).

#### Mixing Score of Binary-categorical Data

To quantize the mixing level of the two types of data (real expression profiles vs. generated expression profiles in our case), we defined a mixing score for a *k*-clustering result of binary-categorical data as follows. Suppose the total number of data to be clustered is *N*. For a cluster *i*, the number of data in this cluster of one category is denoted as *p_i_*, and the number of data of the other category is denoted as *q_i_*. Then the mixing score for a *k*-clustering result is defined as

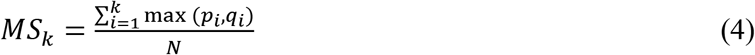

This score equals the average proportions of data from the category that dominates each cluster. The mixing score is of range [0.5, 1], where 0.5 indicates the two categories on average mixing evenly in the *k* clusters, and 1 indicates the two categories are cleanly separated among the *k* clusters. The mixing score tends to increase with the number of clusters *k* used to stratify the data.

### Data and Code Availability

The code generated during this study and pretrained VAE and S-VQ-VAE models are available on GitHub at https://github.com/evasnow1992/DeepGenerativeModelLINCS. This study did not generate datasets.

## Supplemental Information titles and legends

**Figure S1. The architecture of VAE model.** The encoder is composed of three hidden layers, where the third hidden layer defines the distribution from which the encoding vector *z_i_* is sampled. The reparameterization trick is applied to generate a differentiable estimate of *z_i_*, which allows gradient descent to be used for training the model. The decoder has a reversed architecture as the encoder.

**Figure S2. PCA two components scatter plots and density contour plots of three input datasets.** (A and B) Scatter plot and density contour of the SMP dataset. The outlier group on the right of the scatter plot is composed of 4,649 samples treated with bortezomib and MG-132. Both these SMPs are proteasome inhibitors. (C and D) The scatter plot and density contour of the GP dataset. (E and F) The scatter plot and density contour of a combination of the SMP and GP datasets (the SMGP dataset), excluding the outlier group of proteasome inhibitors.

**Table S1. LINCS datasets and major cell lines.** A major cell line is a cell line that has over 10,000 samples.

**Table S2. The data reconstruction performance of VAE models and S-VQ-VAE model on training data and validation data.** The models were trained on 9/10 of the data and validated on the other 1/10 of the data. The reported losses are MSE between the reconstructed data and the input data.

**Table S3. Performance of PCL classification with different sample representations as input data.** The models were trained on 9/10 of the SMC data and tested on the other 1/10. LR: logistic regression; SVM: support vector machine.

**Table S4. The mean of rank of the top known target for 16 FDA-approved drugs from drug-target prediction with different types of representation.** Data related to Figure. 6. The value for a given drug and a type of representation is the mean rank of the drug target gene(s) among all genes when retrieved and ranked according to the similarity between the representation of drug and representations of samples with gene knockdowns. The lowest mean rank is bolded for each drug. # of SMP: the number of SMP samples treated with the drug; # of GP: the number of GP samples with target gene knocked down; E: encoder layer; T: top hidden layer; Sig.: signature nodes; D: decoder layer.

