## Supplementary figures and tables for "Learning to Encode Cellular Responses to Systematic Perturbations with Deep Generative Models"

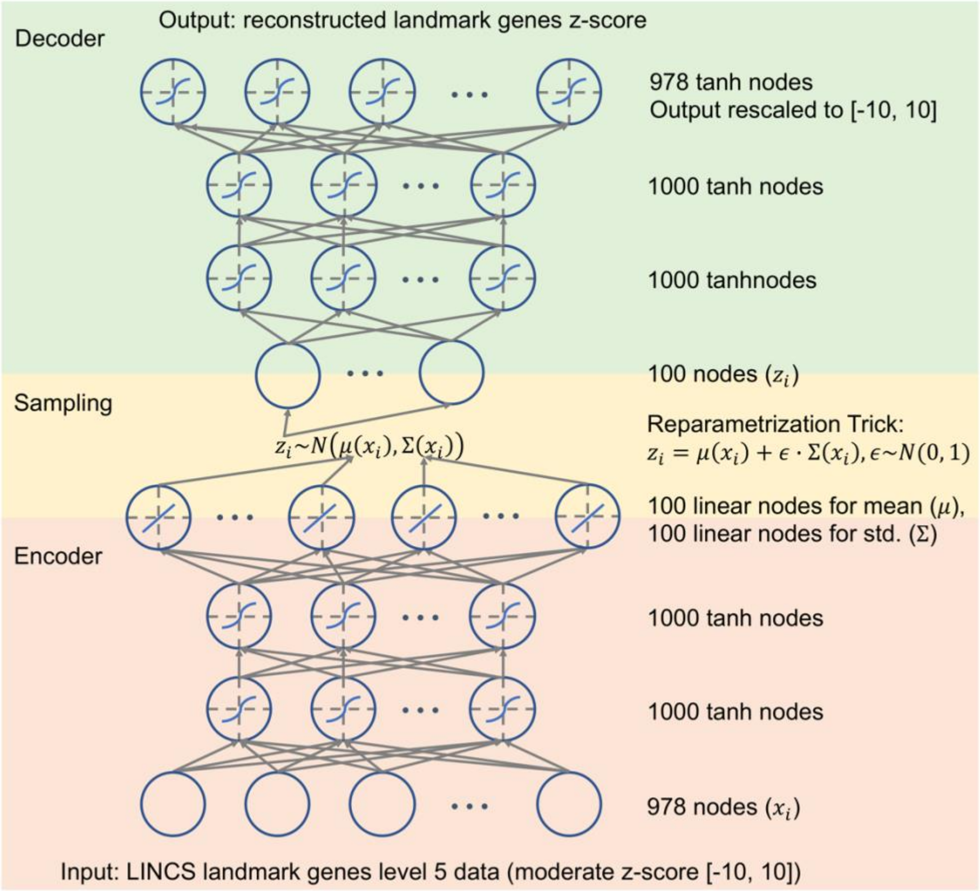


**Figure S1. The architecture of VAE model.** The encoder is composed of three hidden layers, where the third hidden layer defines the distribution from which the encoding vector $z_i$ is sampled. The reparameterization trick is applied to generate a differentiable estimate of $z_i$, which allows gradient descent to be used for training the model. The decoder has a reversed architecture as the encoder.

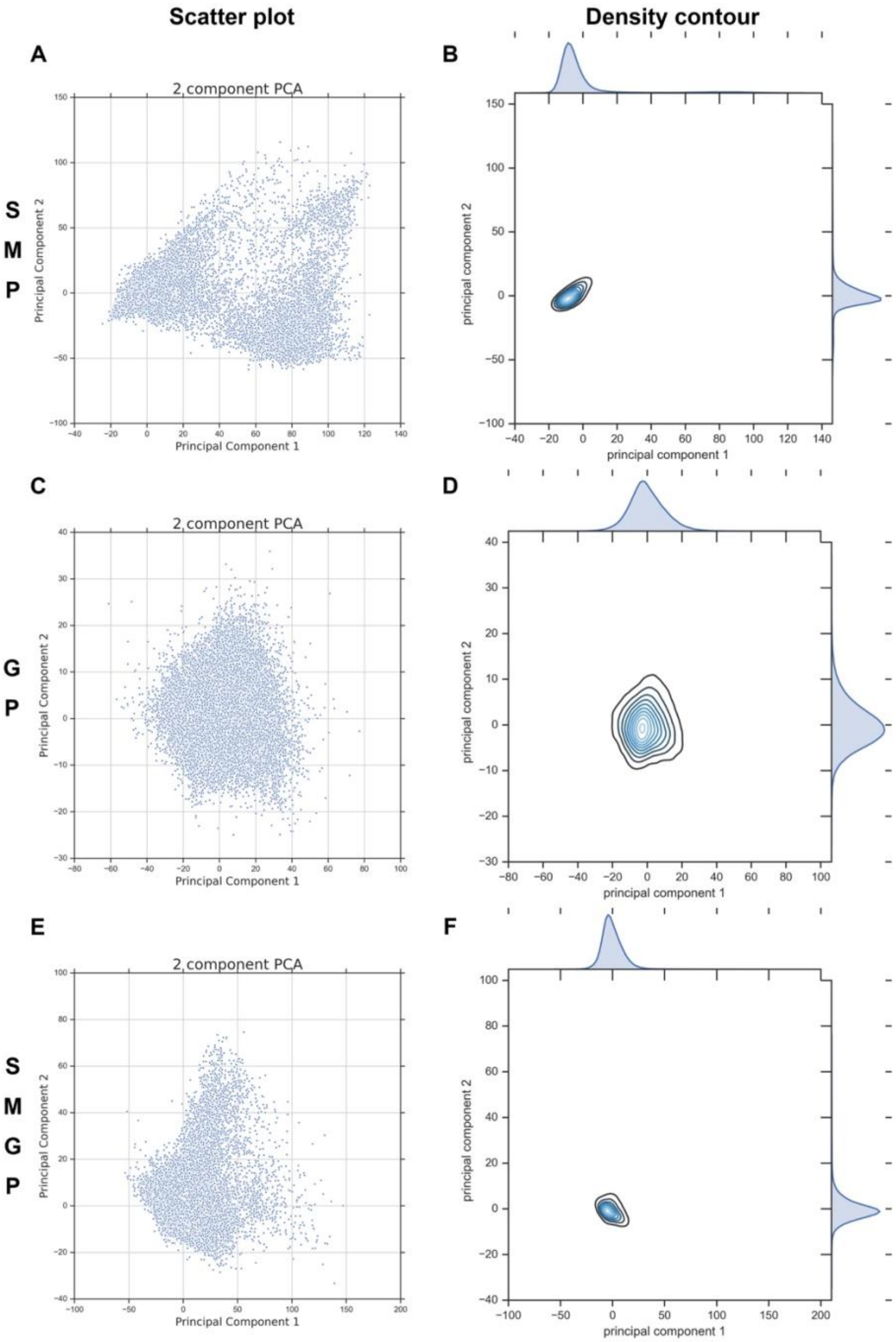


**Figure S2. PCA two components scatter plots and density contour plots of three input datasets.** (A and B) Scatter plot and density contour of the SMP dataset. The outlier group on the right of the scatter plot is composed of 4,649 samples treated with bortezomib and MG-132. Both these SMPs are proteasome inhibitors. (C and D) The scatter plot and density contour of the GP dataset. (E and F) The scatter plot and density contour of a combination of the SMP and GP datasets (the SMGP dataset), excluding the outlier group of proteasome inhibitors.

**Table S1. LINCS datasets and major cell lines.** A major cell line is a cell line that has over 10,000 samples.

| GEO ID | Dataset content | # of sample | # of perturbagen | # of drug/gene name | # of perturbagen class (PCL) | # of cell line | Major cell line name | Major cell line type | # of sample in cell line |
|---|---|---|---|---|---|---|---|---|---|
| GSE70138 | LINCS phase II L1000 dataset, mainly small molecular perturbation | 118050 | 2170 | 1826 | 991 perturbagens in 171 classes | 41 | MCF7 | breast adenocarcinoma | 13476 |
| | | | | | | | A375 | malignant melanoma | 12740 |
| | | | | | | | PC3 | prostate adenocarcinoma | 12719 |
| | | | | | | | HT29 | colorectal carcinoma | 12529 |
| | | | | | | | HA1E | normal kidney | 12481 |
| | | | | | | | YAPC | pancreatic carcinoma | 10621 |
| | | | | | | | HELA | large intestine adenocarcinoma | 10617 |
| GSE106127 | LINCS L1000 RNAi and CRISPR dataset. Corresponds to genetic perturbational signatures of shRNAs and CRISPR reagents, that exist in GEO series GSE70138 and GSE92742. | 119013 | 18413 | 4320 | NA | 15 | VCAP | prostate carcinoma | 17098 |
| | | | | | | | A375 | malignant melanoma | 13121 |
| | | | | | | | PC3 | prostate adenocarcinoma | 13061 |
| | | | | | | | HA1E | normal kidney | 12957 |
| | | | | | | | A549 | non-small cell lung cancer | 12691 |
| | | | | | | | HT29 | colorectal carcinoma | 12305 |
| | | | | | | | HCC515 | lung cancer | 11985 |
| | | | | | | | MCF7 | breast adenocarcinoma | 11869 |
| | | | | | | | HEPG2 | hepatocellular carcinoma | 11695 |

**Table S2. The data reconstruction performance of VAE models and S-VQ-VAE model on training data and validation data.** The models were trained on 9/10 of the data and validated on the other 1/10 of the data. The reported losses are MSE between the reconstructed data and the input data.

| Model type | Train data | Training loss | Validation loss |
|---|---|---|---|
| VAE | SMP | 1.081 | 1.113 |
| VAE | GP | 0.861 | 0.864 |
| VAE | SMGP | 0.995 | 1.002 |
| S-VQ-VAE | SMCNP | 1.725 | 1.831 |

**Table S3. Performance of PCL classification with different sample representations as input data.** The models were trained on 9/10 of the SMC data and tested on the other 1/10. LR: logistic regression; SVM: support vector machine.

| Representation type | LR test accuracy | SVM test accuracy |
|---|---|---|
| raw | 0.5922 | 0.5273 |
| encoder layer 1 | 0.5096 | 0.4528 |
| encoder layer 2 | 0.4461 | 0.3881 |
| encoder layer 3 | 0.4098 | 0.4232 |
| signature nodes | 0.3814 | 0.3615 |
| decoder layer 1 | 0.4002 | 0.4082 |
| decoder layer 2 | 0.3994 | 0.4085 |

**Table S4. The mean of rank of the top known target for 16 FDA-approved drugs from drug-target prediction with different types of representation.** Data related to Figure. 6. The value for a given drug and a type of representation is the mean rank of the drug target gene(s) among all genes when retrieved and ranked according to the similarity between the representation of drug and representations of samples with gene knockdowns. The lowest mean rank is bolded for each drug. # of SMP: the number of SMP samples treated with the drug; # of GP: the number of GP samples with target gene knocked down; E: encoder layer; T: top hidden layer; Sig.: signature nodes; D: decoder layer.

| Drug | Target | # of SMP | # of GP | Raw | E1 | E2 | T | Sig. | D1 | D2 |
|---|---|---|---|---|---|---|---|---|---|---|
| pitavastatin | HMGCR | 42 | 45 | **376.40** | 962.69 | 1930.62 | 806.81 | 706.74 | 827.60 | 688.00 |
| bortezomib | PSMB10, PSMA3, PSMA1, PSMA5, PSMB7, PSMB5, PSMA8, PSMB1 | 2339 | 196 | 20.32 | 48.12 | 45.94 | 65.69 | 96.78 | 49.09 | **19.13** |
| hydrocortisone | NR3C1 | 42 | 36 | 4605.67 | 3759.6 | 2221.74 | 3019.07 | 3339.52 | **2122.33** | 2392.12 |
| vemurafenib | BRAF | 81 | 108 | 1030.01 | 981.65 | 1215.56 | 779.80 | **664.57** | 970.10 | 1060.69 |
| flutamide | AR | 42 | 29 | **5111.36** | 7178.43 | 8167.55 | 9172.00 | 11013.07 | 13934.31 | 13047.52 |
| clobetasol | NR3C1 | 42 | 36 | 5221.50 | 5025.12 | 3236.19 | 5060.95 | 5628.48 | **3136.02** | 4185.90 |
| digoxin | ATP1A3, FXYD2, ATP1B1 | 42 | 83 | 595.62 | 431.10 | 492.21 | 497.86 | **336.48** | 590.71 | 405.69 |
| mycophenolate-mofetil | IMPDH2 | 42 | 27 | **4594.12** | 6514.81 | 5928.67 | 6630.00 | 5091.55 | 8631.02 | 4683.26 |
| dasatinib | LCK, YES1 | 204 | 175 | 942.70 | 743.19 | 694.73 | 681.38 | **641.82** | 649.23 | 658.83 |
| amlodipine | CACNA1D | 42 | 27 | 7798.79 | 4797.38 | 5290.60 | 5252.71 | 5159.55 | 5872.90 | **4295.33** |
| calcitriol | VDR | 42 | 27 | 5293.00 | 5859.21 | 6924.50 | 5877.14 | 6128.60 | **5263.52** | 5856.98 |
| glibenclamide | KCNJ11 | 42 | 27 | 6074.93 | 9643.98 | 8653.05 | 7038.45 | **3969.43** | 7179.50 | 7104.81 |
| paclitaxel | TUBB6, TUBA1A, TUBB2A, TUBB2C | 42 | 105 | 464.17 | 548.00 | 781.81 | 640.90 | **412.62** | 421.00 | 831.40 |
| losartan | AGTR1 | 42 | 24 | 4841.31 | 4767.88 | 4162.24 | 3865.98 | 3871.10 | **3656.90** | 4469.76 |
| irinotecan | TOP1 | 42 | 27 | 5079.93 | 5317.57 | 4332.83 | 4459.55 | 4248.19 | 3999.52 | **3915.17** |
| raloxifene | ESR2 | 42 | 36 | 4686.60 | 3236.07 | **1262.83** | 2078.57 | 2383.14 | 1595.02 | 1757.98 |
